# Decoding the Effects of Spike Receptor Binding Domain Mutations on Antibody Escape Abilities of Omicron Variants

**DOI:** 10.1101/2022.07.21.500931

**Authors:** Sandipan Chakraborty, Aditi Saha, Chiranjeet Saha, Sanjana Ghosh, Trisha Mondal

**Affiliations:** Center for Innovation in Molecular and Pharmaceutical Sciences (CIMPS), Dr. Reddy’s Institute of Life Sciences, University of Hyderabad Campus, Gachibowli, Hyderabad 500046, India; Amity Institute of Biotechnology, Amity University, Kolkata, 700135, India

**Keywords:** Omicron variants, Mutations, Antibody escape, Immunogenic hotspot, Binding affinity

## Abstract

Recent times witnessed an upsurge in the number of COVID cases which is primarily attributed to the emergence of several omicron variants, although there is substantial population vaccination coverage across the globe. Currently, many therapeutic antibodies have been approved for emergency usage. The present study critically evaluates the effect of mutations observed in several omicron variants on the binding affinities of different classes of RBD-specific antibodies using a combined approach of immunoinformatics and binding free energy calculations. Our binding affinity data clearly show that omicron variants achieve antibody escape abilities by incorporating mutations at the immunogenic hotspot residues for each specific class of antibody. K417N and Y505H point mutations are primarily accountable for the loss of class I antibody binding affinities. The K417N/Q493R/Q498R/Y505H combined mutant significantly reduces binding affinities for all the class I antibodies. E484A single mutation, on the other hand, drastically reduces binding affinities for most of the class II antibodies. E484A and E484A/Q493R double mutations cause a 33-38% reduction in binding affinity for the approved therapeutic monoclonal antibodies, Bamlanivimab (LY-CoV555). The Q498R RBD mutation observed across all the omicron variants can reduce ~12% binding affinity for REGN10987, a class III therapeutic antibody, and the L452R/Q498R double mutation causes a ~6% decrease in binding affinities for another class III therapeutic antibody, LY-CoV1404. Our data suggest that achieving the immune evasion abilities appears to be the selection pressure behind the emergence of omicron variants.

## 1. Introduction

The recent emergence of a new beta-coronavirus, Severe Acute Respiratory Syndrome Coronavirus 2 (SARS-CoV2), has led to the ongoing pandemic, which completely disrupted the public health care system and caused mortality of more than 6 million people. The taxonomic classification reveals that the virus belongs to the order Nidovirales, family Coronaviridae, subfamily Orthocoronavirinae, genus Betacoronavirus, subgenus Sarbecovirus, and realm Riboviria.^1^ The genetic material, a positive sense RNA genome wrapped around the nucleocapsid protein, is enclosed within a lipid envelope embedded with three structural proteins: the spike glycoprotein (S), an envelope protein (E), and membrane (M).^2^

The spike protein contains many immunogenic regions^3–5^ and mediates the recognition of the human angiotensin converting enzyme 2 (ACE2) receptor.^6^ As a result, the spike protein is the prime target for vaccine development and neutralizing antibody development.^7–11^ Most neutralizing antibodies target the receptor-binding domain (RBD) or the N-terminal domain (NTD) of the spike. Notably, the spike monomer consists of the S1 and S2 domains. Both the NTD and RBD exist within the S1 domain.^12,13^ Spike monomer entangles to form spike trimer. Each RBD shows exquisite conformational plasticity within the spike trimer and exists either in the “Up” or “Down” conformation.^14,15^ The “Up” RBD is capable of successful ACE2 binding.

Food and Drug Administration (FDA) approved several anti-SARS-CoV2 mAbs for emergency usage. The combination of bamlanivimab (LY-CoV555) and etesevimab neutralizing mAbs has been approved to treat mild to moderate COVID-19 with a high risk of hospitalization.^16^ The mAbs bind to the overlapping epitopes on spike RBD. Bebtelovimab (LY -CoV1404) is a recombinant anti-RBD neutralizing mAb.^17^ Casirivimab and imdevimab (REGN-COV) recombinant anti-RBD mAbs bind to the non-overlapping epitopes.^18^ Sotrovimab also received emergency approval for mild to moderate COVID19.^19^ It is effective against both SARS-CoV and SARS-CoV2, as it recognizes the common epitopes in both the RBDs. The combination of anti-RBD mAbs, tixagevimab (AZD8895)/cilgavimab (AZD1061), also received emergency approval from FDA.^20^ As the pandemic went on, new variants emerged.^21–24^ Alpha, beta, gamma, delta, and omicron are the variants of concerns (VOCs) that caused a massive surge of cases of infection and mortality across the globe. Besides, many variants of interest (VOIs) cause local epidemics and are under strict surveillance (https://viralzone.expasy.org/9556). Different omicron variants primarily contribute to the recent increase in COVID-19 case fatality.^25^ Efficacy of neutralizing antibodies and vaccines against these variants is still highly doubtful.^26–29^ Bamlanivimab and etesevimab combination therapy and sotrovimab are not beneficial against omicron variants.^30^ Therefore, their usage has been stopped in the USA. Bebtelovimab showed efficacy against all circulating omicron subvariants *in vitro*,^1^ but its effectiveness in clinical settings is yet to be proven. Apart from this, many therapeutic antibodies showed efficacy *in vitro*. B38,^31^ CA1,^32^ CB6,^32^ CR3022,^33^ S309,^34^ BD368-2,^35^ and many more are anti-SARS-CoV2 RBD therapeutic antibodies demonstrated efficacy against SARS-CoV2.^36^

RBD-specific neutralizing Abs are classified as class I, II, III, and IV.^37^ Class I Abs bind RBD in the “up” conformation at the receptor-binding motif (RBM), thereby impeding ACE2 binding. Class II mAbs, on the other hand, bind RBD in both up and down conformation at the RBM. Both the class I and class II Abs are “ACE2-blocker”. Class III NAbs bind outside the RBM and recognizes both the “up” and “down” conformation. The class IV Abs do not overlap with the ACE2 binding site and bind to the conserved cryptic epitope at the base of the RBD.

Data analysis during the early pandemic periods reveals that the critical mutations appear at the RBD that enhance ACE2 recognition.^22,38^ Thus better receptor usage is the driving force for selecting SARS-CoV2 RBD variants. Later, detailed molecular dynamics simulations revealed that the N501Y and E484K mutations observed in different VOCs enhance ACE2 recognition but reduce antibody binding.^39^ Many mutations in the RBD regions are shown to escape antibody recognition. K417T,^40^ N439K,^41^ G446V,^42^ N450K,^43^ A475V,^40^ T478I,^43^ F490S,^21^ and S494P^44^ show antibody escape capability. Thus, it is essential to analyze the spike-antibody interactions to elucidate the critical epitope-paratope interactions that enhance our understanding of the antibody escape abilities of rapidly circulating VOCs.

The last two years witnessed rapid growth in spike-antibody crystal structures due to the unmet need of the time. All the spike-antibody structures are curated in the CoV3D database.^45^ The present study critically explores all spike-antibody complexes from different classes using protein-protein interaction analysis to explore critical epitope-paratope interactions responsible for different classes of antibody recognition. The binding mode and epitopes for class IV antibodies are very different and do not interfere with ACE2 binding. Therefore, we have not considered class IV antibodies for this study. Also, the possibility of immune escape as a selection pressure on currently circulating SARS-CoV2 variants has been explored using several immunoinformatics analysis. Finally, the RBM mutations that appear at critical hotspot residues for each antibody class have been identified for all the omicron variants, and the effect of these mutations on the antibody binding affinities have been calculated.

## 2. Materials and Methods

### 2.1. Interaction analysis of Spike-antibody complexes

Experimentally resolved structures of all the available spike-antibody complexes belonging to four different classes were obtained from the CoV3D database (https://cov3d.ibbr.umd.edu/antibody_classification).^45,46^ The dataset contained spike-antibody complexes for 41 class I antibodies, 69 class II antibodies, and 32 class III antibodies. Spike-antibody interactions were analyzed with the Protein Interaction Calculator (PIC) webserver (http://pic.mbu.iisc.ernet.in/).^47^

### 2.2. Immunoinformatics analysis

Epitope prediction was carried out using the Immune Epitope Database and Analysis Resource (IEDB). The sequence of the RBD region of the SARS-CoV2 spike protein sequence (319-541) was obtained from the NCBI protein sequence database and used for IEDB analysis (http://www.iedb.org/).^48^ Antibody epitope prediction and physicochemical characterization of the epitopes were carried out using the Bepipred Linear Epitope Prediction 2.0, Chou and Fasman Beta-Turn Prediction, Emini Surface Accessibility Prediction, Karplus and Schulz Flexibility Prediction, Kolaskar and Tongaonkar Antigenicity and Parker Hydrophilicity Prediction tools.

### 2.3. Effect of spike RBD mutations observed in the omicron variants on different classes of antibody recognition

The curated crystal structures of eight class I, nine class II and four class III SARS-CoV2 RBD-antibody complexes were obtained from the CoV3D database. We considered four single RBD mutants and one combined RBD mutant for class I antibody complexes. These were K417N, Y505H, Q493R, Q498R single RBM mutants, and the combined mutant contained all four mutations (K417N, Y505H, Q493R, and Q498R). We considered the E484A and Q493R single RBD mutants and a dual mutant containing both the mutations for class II antibodies. L452R and Q498R single mutants and a dual mutant containing both the mutations were considered for class III antibodies. Mutant RBD-antibody complexes were generated using computational mutagenesis. The binding free energies of wild-type and mutant RBDs with each antibody were calculated using the MM/GBSA method implemented in the HawkDock web server.^49^

## 3. Results and Discussions

### 3.1. Interaction feature of class I antibody recognition

We have analyzed the crystal structure of forty one class I spike-antibody complexes, and the interaction heatmap is shown in Figure 1. Interactions are deconvoluted in hydrogen bonds, hydrophobic interactions, ionic interactions, π-π stacking interactions, and π-cation interactions. Many RBD discontinuous epitopes are evident from the heatmap for this class of antibodies. Apparent from the heatmap, the Tyr505 is a critical spike RBD residue that makes hydrogen bonding interactions with almost all the class I antibodies. However, the number of hydrogen bonds varies significantly among different antibodies. Apart from that, Tyr489, Asn487, Ala475, Tyr473, Arg457, Leu455, Tyr421, Asp420, Lys417 and Arg403 residues form several hydrogen-bonding contacts with almost all the class I antibodies. Tyr505, Tyr489, Phe486, Phe456, Leu455 and Tyr421 form conserved hydrophobic interactions with all the class I antibodies. Lys417 is the only residue interacting with almost all the class I antibodies through salt-bridge interactions. Tyr505, Phe456, and to some extent Tyr421 form conserved π-π stacking interactions with this class of antibodies. On the other hand, Tyr489 and Phe486 are involved in π-cation interactions with most of the antibodies, while Lys417 forms π-cation interactions with more than 50% of class I antibodies studied here.

**Figure 1:**
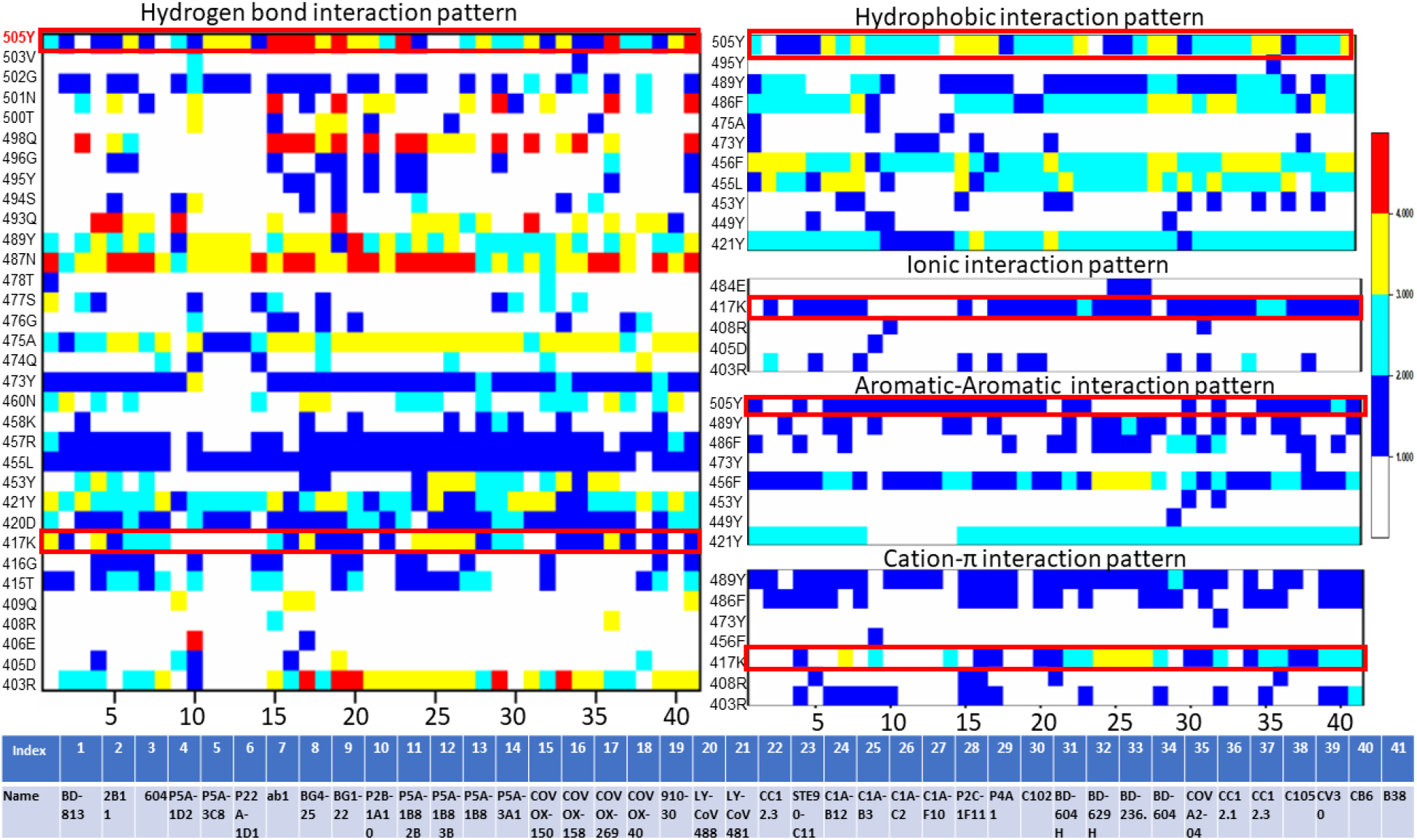
Interaction heatmaps of 41 SARS-CoV2 RBD-class I complex structures. RBD residues form hydrogen bonds, hydrophobic interactions, ionic interactions, π-π interactions, and π-cation interactions with different class I antibodies are shown. Each interaction is colored based on the number of occurrences. Details of the antibodies are also tabulated. Mutations observed in omicron variants at a particular position are shown within a red box.

Few critical hotspot residues form many conserved interactions with almost all the class I antibodies. Tyr505 has involved in hydrogen bonds, hydrophobic and π-π stacking interactions with almost every class 1 antibody. Likewise, Lys417 is another residue involved in conserved hydrogen-bonding interactions, salt-bridge, and π-cation with the antibodies. Interestingly, mutations have been observed in both the residues in the omicron variants. In Omicron variants, Tyr505 and Lys417 have been mutated to Histidine and Asparagine, respectively (marked within a red square in Figure 1). These radical substitutions are expected to reduce antibody recognition for this class.

Tyr489 is also involved in conserved hydrogen bonding, hydrophobic, π-π stacking and π-cation interactions. Similarly, Phe486 is also participating in conserved hydrophobic, π-π stacking, and π-cation interactions with almost all the antibodies. Previously, using MM/GBSA binding free energy calculations, it has been shown that these two residues are the most important ones during ACE2 recognition.^22^ Thus, involving these residues in antibody binding outcompetes ACE2 recognition.

### 3.2. Interaction feature of class II antibody recognition

We have analyzed 69 SARS-CoV2 spike RBD-class II antibody complexes using protein-protein interaction analysis to depict the conformational epitopes on the RBD responsible for class II antibody recognition. Interactions have been decoded in terms of hydrogen bonding, hydrophobic and ionic interactions. Unlike class I, conserved epitopes are less evident for class II antibodies. The two most conserved RBD residues that form hydrogen bonding contacts with most of the class II antibodies are Gln493 and Glu484 (within the red square in Figure 2). Interestingly, mutations have been observed in both the residues in the omicron variants.

**Figure 2:**
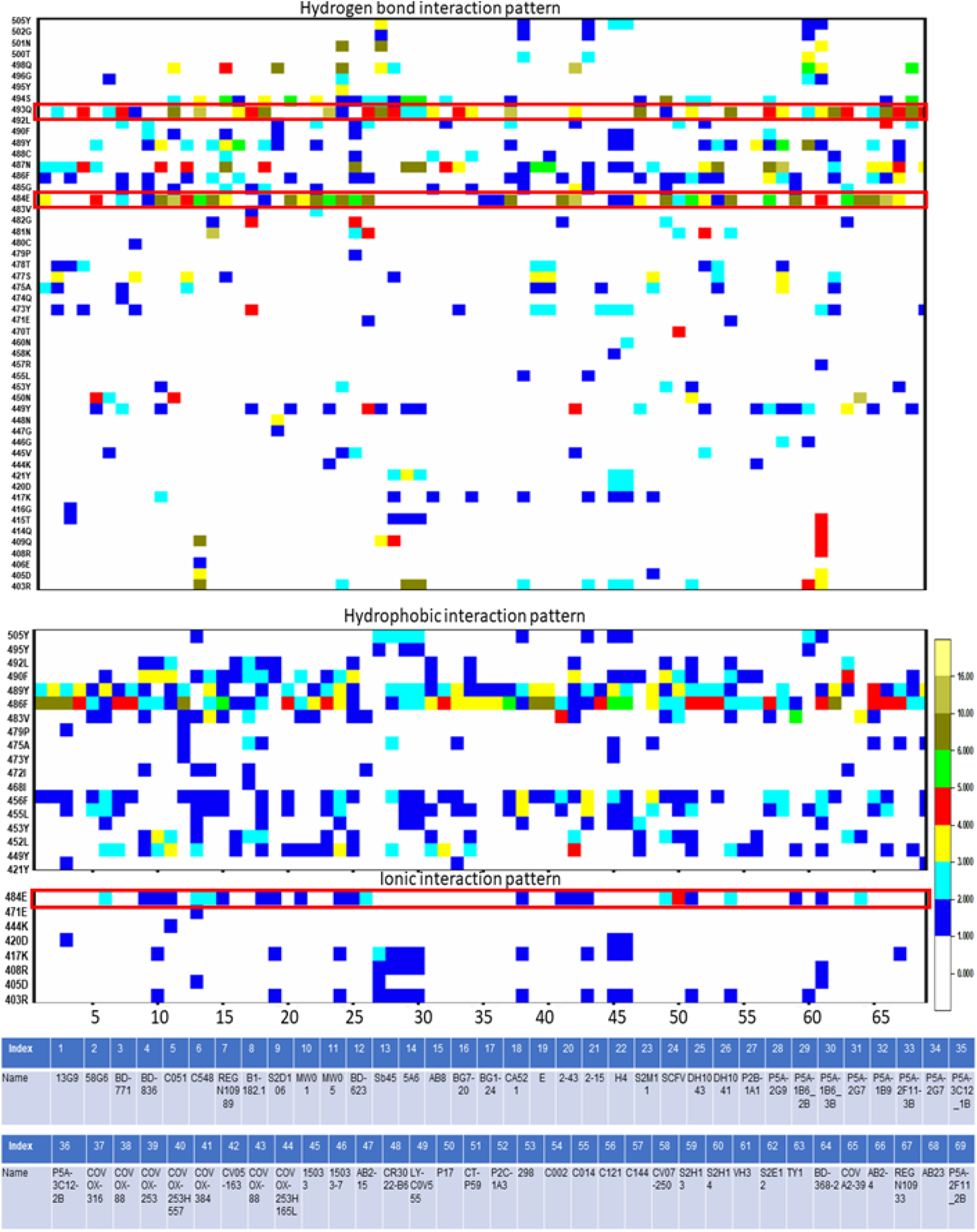
Interaction heatmaps of 69 SARS-CoV2 RBD-class II antibody complex structures. RBD residues form hydrogen bonds, hydrophobic interactions, and ionic interactions with different class II antibodies are shown. Each interaction is colored according to the number of occurrences. Details of the antibodies are also tabulated. Mutations observed in omicron variants at a particular position are shown within a red box

**Figure 3:**
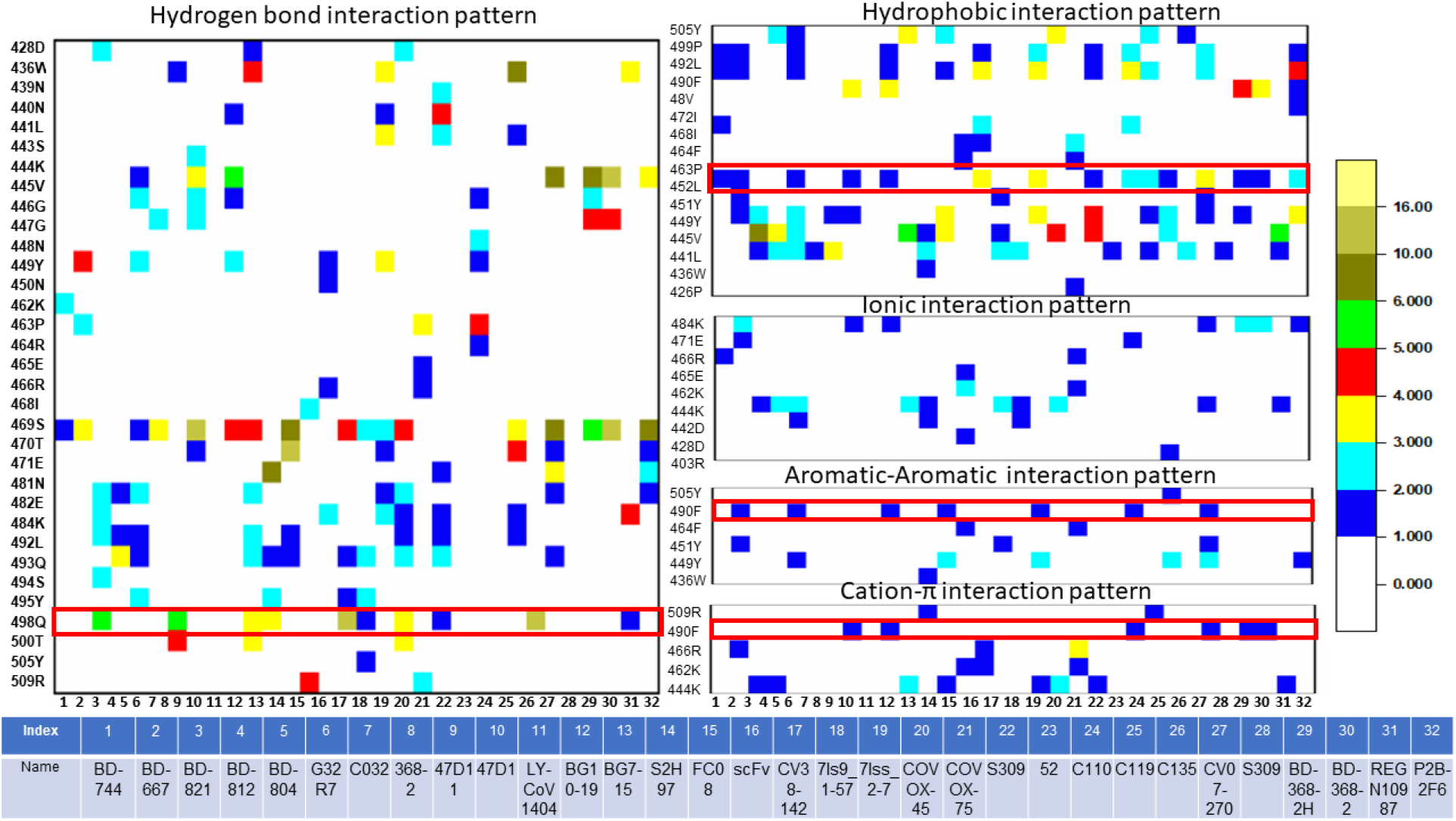
Interaction heatmaps of 32 SARS-CoV2 RBD-class III complex structures. RBD residues form hydrogen bonds, hydrophobic interactions, ionic interactions, π-π interactions, and π-cation interactions with different class III antibodies are shown. Each interaction is colored according to the number of occurrences. Details of the antibodies are also tabulated.

In omicron, the 493^rd^ residue has been mutated to arginine, and the 484^th^ residue has been mutated to alanine. Again both the mutations are very drastic and expected to hinder antibody recognition. Glu484 is also involved in ionic interactions with many class II antibodies.

Tyr489 and Phe486 show highly conserved hydrophobic interactions with almost all the class II antibodies. Notably, these two residues exhibit the highest contribution in binding free energy during ACE2 binding.^22^ Thus the involvement of these residues in the class II antibody binding makes these antibodies competitive with ACE2 binding.

### 3.3. Interaction feature of class III antibody recognition

Meta-analysis of 32 RBD-class III antibody complexes reveals that the epitopes for class III antibodies are highly dispersed. Ser469 appears to be a conserved residue involved in hydrogen bonding interactions with most of the antibodies from this class. Gln498 is also involved in hydrogen bonding interactions with many antibodies from this class. Notably, this residue has mutated to Arg498 in omicron variants.

Leu452 is a crucial RBD epitope to recognize class III antibodies. It involves hydrophobic interactions with almost ~50% of the antibodies studied here. In delta, lambda (C.37+C.37.1), epsilon (B. 1.427, B.1.429) and Kappa (B.1.671.1) variants contain mutation on the particular residues (L452R). Also, the recent omicron subvariants BA.4 and BA.5 contain this particular mutation which can be interpreted by the antibody escape abilities of those variants. On the other hand, Phe490 shows π-π stacking and π-cation interactions with few of the antibodies. This residue has been observed to mutate to Ser490 in lambda variants.

### 3.4. Understanding the immunogenic potential of RBM hotspot residues

The physicochemical properties behind the epitopic potential of the identified hotspot RBM residues have been decoded using various immunoinformatics algorithms. We have considered the highly conserved hotspot residues of spike RBM responsible for class I, II, and III antibody recognition and analyzed their antigenicity, surface accessibility, hydrophilicity, beta turns, and flexibility using the IEDB webserver.^48^ Results are summarised in Table 1.

**Table 1:** Immunogenic properties of the identified SARS-CoV2 RBM hotspot residues

| PROPERTIES | RBM RESIDUE NUMBER |  |  |  |  |  |  |
| --- | --- | --- | --- | --- | --- | --- | --- |
|  | *Lys417* | *Leu452* | *Glu484* | *Gln493* | *Phe490* | *Gln498* | *Tyr505* |
| **ANTIGENICITY** | 0.973 | **1.090** | 1.037 | 1.067 | **1.110** | 0.984 | **1.076** |
| **SURFACE ACCESSIBILITY** | 0.879 | **1.369** | 0.247 | 1.195 | 0.418 | **1.386** | **1.257** |
| **HYDROPHILICITY** | **3.814** | **1.843** | **2.100** | 0.000 | 0.543 | **2.129** | **1.714** |
| **BETA TURNS** | 1.014 | 1.016 | 1.101 | **1.117** | 1.083 | **1.189** | 1.200 |
| **FLEXIBILITY** | **1.035** | 0.928 | 0.988 | 0.986 | 0.913 | 1.009 | 0.990 |

responsible for antibody recognition

Lys417 is highly hydrophilic and possesses high flexibility and ß-turn propensity, which are the two dominant characteristics of immunogenic epitopes. Leu452, Phe490, and Tyr505 have been predicted to be immunogenic. Leu452, Gln498, and Tyr505 are highly surface exposed, and Gln498 and Tyr505 are highly hydrophilic. Thus they are accessible for antibody recognition. Glu484 is buried, but its high hydrophilicity makes it appropriate for class II antibody recognition. Gln493 shows moderate antigenicity, surface accessibility, and β-turn propensity making it suitable to recognize the class I and II antibodies. Phe490 has been predicted to be highly antigenic.

Interestingly, many of these residues involved in different antibody recognition classes get mutations in circulating omicron variants. A list of RBM mutations observed in different omicron lineages^50^ is summarized in Table 2.

**Table 2:** List of RBD mutations observed in several omicron variants. Mutations responsible for different classes of antibody recognition are coloured differently.

| Omicron variants |  |  |  |  |  |
| --- | --- | --- | --- | --- | --- |
| BA.1 | BA.2 | BA.2.12.1 | BA.3 | BA.4 | BA.5 |
| G339D | G339D | G339D | G339D | G339D | G339D |
| S371L | S371F | S371F | S371F | S371F | S371F |
| S373P | S373P | S373P | S373P | S373P | S373P |
| S375F | S375F | S375F | S375F | S375F | S375F |
| K417N | T376A | T376A | D405N | T376A | T376A |
| N440K | D405N | D405N | K417N | D405N | D405N |
| G446S | R408S | R408S | N440K | R408S | R408S |
| S477N | K417N | K417N | G446S | K417N | K417N |
| T478K | N440K | N440K | S477N | N440K | N440K |
| E484A | S477N | L452Q | T478K | L452R | L452R |
| Q493R | T478K | S477N | E484A | S477N | S477N |
| G496S | E484A | T478K | Q493R | T478K | T478K |
| Q498R | Q493R | E484A | Q498R | E484A | E484A |
| N501Y | Q498R | Q493R | N501Y | F486V | F486V |
| Y505H | N501Y | Q498R | Y505H | Q498R | Q498R |
|  | Y505H | N501Y |  | N501Y | N501Y |
|  |  | **Y505H** |  | **Y505H** | **Y505H** |

*Red: Class I; Light blue: Class II; Orange: Class III*

The table shows that K417H and Y505H mutations are common to all lineages of omicron subvariants. These two residues are involved in almost all the class I antibody recognition. 493^rd^ and 484^th^ residues are essential for class II antibody recognition. All omicron sublineages contain E484A, and the Q493R mutation has been observed in BA.1, BA.2, BA.2.12.1, and BA.3 variants. Q498R mutation has been observed in all omicron subvariants, but L452R mutations have been observed in only BA.4 and BA.5 sub-variants. Both these residues are specific hotspots for class III antibody binding.

### 3.5. Understanding the effect of RBM mutations present in omicron variants on different classes of antibody recognition: Binding free energy analysis

Class I and II antibodies primarily bind to the RBM and outcompete ACE2 binding. Although class III antibodies do not directly compete with ACE2 for the same binding site, their binding to other parts of RBD sterically hinders ACE2 recognition. We have considered the class I, II and III antibodies for binding free energy calculations to understand the effect of individual RBM mutations observed in omicron variants on antibody recognition. Mutants are generated at a particular residue using computational mutagenesis, and binding free energies were calculated using MM/GBSA method implemented in the HawkDock webserver.^49^

#### 3.5.1. Effect of RBM mutations at the hotspot residues observed in omicron variants on class I antibody recognition

As evident from the interaction heatmap for class I antibodies, two highly conserved interacting residues are Lys417 and Tyr505. In all omicron variants, we observed the appearance of K417N and Y505H mutations. Since Gln493 and Gln498 are also involved in critical interactions with many antibodies, we have also considered the Q493R and Q498R RBM mutations observed in omicron variants. We have analyzed the effect of these single RBD mutations on the binding free energies with eight different class I antibodies. These antibodies are CC12.1, Covox-150, BD-604, P5A-3C8, P22A-1D1, C1A-B12, C1A-F10 and 910-30. We choose these antibodies because all the hotspot amino acids are involved in one or many interactions with these antibodies. We have also considered a combined mutant where all four mutations (K417N, Y505H, Q493R, and Q498R) have been incorporated within the RBD, as observed in omicron variants, to understand the class I antibody escape abilities of omicron variants.

As evident from Figure 4, K419N mutation alone significantly impacts antibody binding. Apart from BD-604, there is a substantial reduction in binding affinity for all other studied antibodies. Notably, K417N RBD mutation causes a ~ 5-10% reduction in binding affinities for COVOX-150, P5A-3C8, C1A-B12, and C1A-F10 antibodies. Evident effects have also been observed for Y505H mutation, which causes a modest decrease in binding affinity for CC12.1, BD-604, P5A-3C8, P22A-1D1, C1A-F10, and 910-30. On the other hand, Q493R and Q498R mutations do not impact class I antibody recognition. However, in the case of the combined mutant, a significant reduction in binding affinities has been observed for all the studied class I antibodies (~20% reduction in binding affinities). These data suggest that omicron variants can reduce the neutralization abilities of class I antibodies.

**Figure 4:**
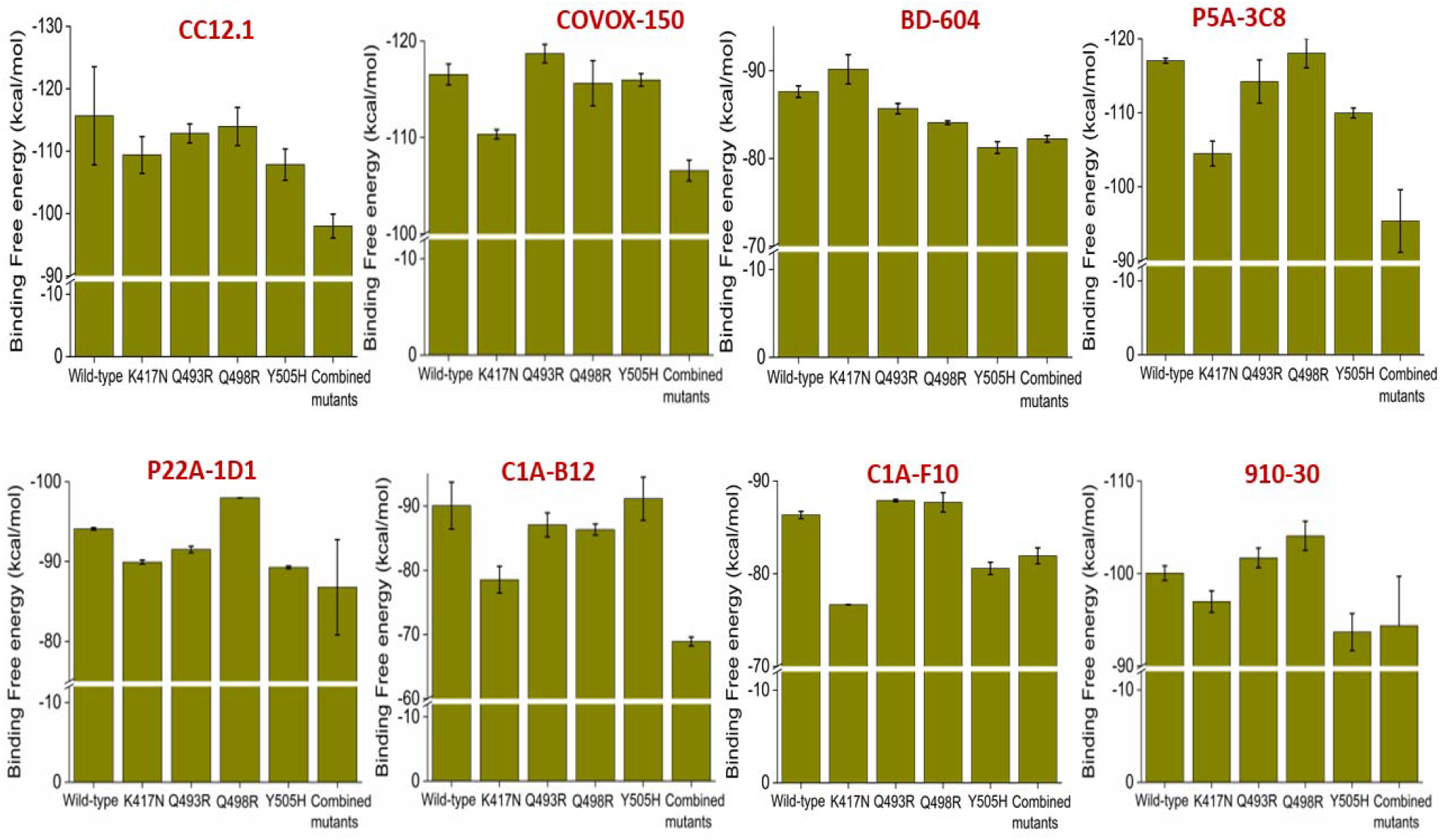
Effect of K417N, Q493R, Q498R, Y505H single SARS-CoV2 RBD mutations and K417N/Q493R/Q498R/Y505H combined SARS-CoV2 RBD mutations on the binding affinities for eight different class I antibodies are shown. Values are represented as Mean ± S.D. (n=3).

#### 3.6.2. Effect of RBM mutations at the hotspot residues observed in omicron variants on class II antibody recognition

Omicron variants contain two RBM mutations (E484A and Q493R) at the hotspot regions specific for class II antibody recognition. The effect of these RBM mutations to the binding abilities of nine different class II antibodies have been studied using the binding free energy calculations. We have considered scFv, MW05, CA521, VH3, C002, LY-CoV555, DH1043, CV05-163, and Sb45, since both the RBD hotspot residues (484^th^ and 493^rd^) are involved in one or many interactions with all these antibodies. We have also generated a double mutant where both the mutations are embedded within the RBD to understand the class II antibody escape abilities of omicron variants. Results are summarized in Figure 5.

**Figure 5:**
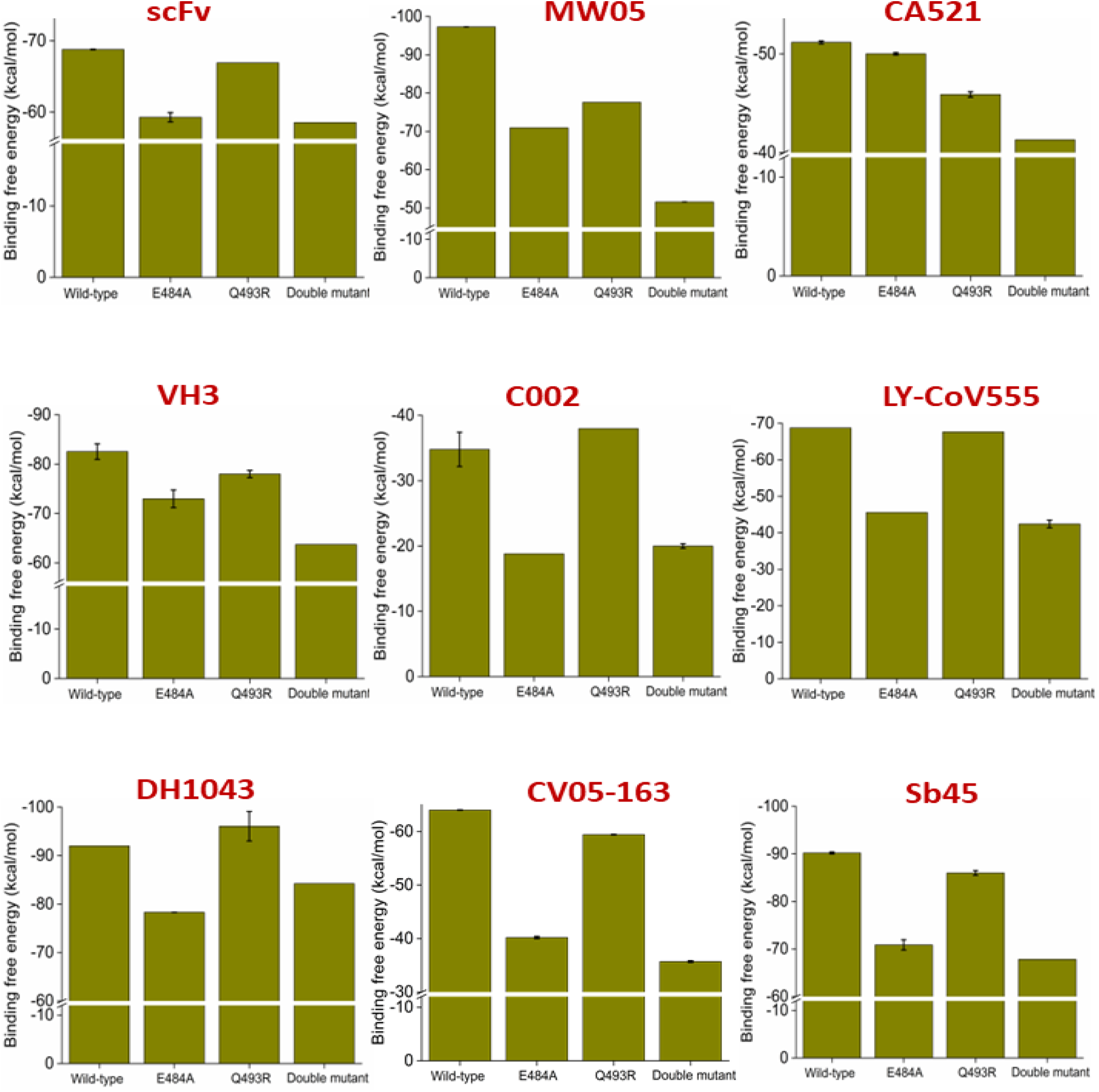
Effect of E484A, Q493R single SARS-CoV2 RBD mutations and E484A/Q493R dual RBD mutation on the binding affinities for nine different class II antibodies are shown. Values are represented as Mean ± S.D. (n=3).

Our data suggest a dramatic decrease in antibody binding affinity. The E484A mutant can cause a ~20% reduction in class II antibody binding affinities, except for CA-521. On the other hand, Q493R mutation causes a substantial decrease in binding affinity for that particular antibody. Apart from CA-521, Q493R mutation reduces the binding affinity for MW05 only. However, the double mutant significantly reduces binding affinities for all the studied class II antibodies. We observe a ~45% decrease in binding affinities for the double mutant for the MW05, C002, and CV05-163. Evident from Table 2 that the BA.1, BA.2, BA.2.12.1, and BA.3 omicron variants contains both mutations. Therefore, they are capable of escaping class II antibody-mediated neutralization. Notably, both the E484A and the double mutations observed in omicron variants significantly reduce (33-38%) the binding affinity for Eli Lilly’s neutralizing monoclonal antibodies, Bamlanivimab (LY-CoV555).

#### 3.5.2. Effect of RBM mutations at the hotspot residues observed in omicron variants on class III antibody recognition

Although class III antibodies do not directly bind to RBM, they sterically interfere with ACE2 binding with the RBM. Therefore, we looked into the interaction heatmap and identified two moderately conserved residues, Leu452 and Gln498. All the omicron variants contain a Q498R RBM mutation, and recently emerged BA.4 and BA.5 variants contain another mutation, L452R. We have studied the effect of both the mutation individually and in combination on four class III antibody recognitions. Results are summarized in Figure 6.

**Figure 6:**
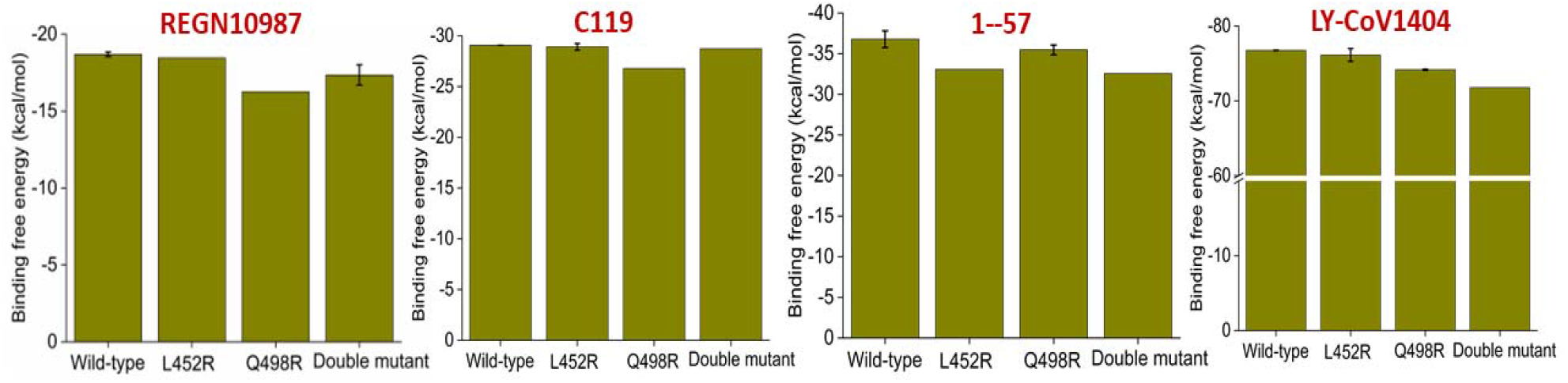
Effect of L452R, Q498R single SARS-CoV2 RBD mutations, and L452R/Q498R dual RBD mutation on the binding affinities for four different class III antibodies are shown. Values are represented as Mean ± S.D. (n=3).

The Q498R mutation causes a moderate decrease in binding affinities for REGN10987, and C119, while the L452R mutation reduces a borderline reduction in binding affinities for 1-57.

The double mutation causes ~6% decrease in binding affinities for the LY-CoV1404 (Eli Lilly’s IgG1 MAbs). This observation is expected, as we have considered the RBM mutations only in circulating omicron variants while this class of antibodies does not directly bind to the RBM.

## 4. Conclusions

The present study demonstrates that omicron variants achieve mutations at critical hotspot residues involved in recognition of different classes of antibodies. Recent simulation and experimental studies indicate that omicron shows comparable binding affinities for ACE2 like the wild-type RBD and a much weaker binding affinity for ACE2 than the delta variants.^51^ Thus better receptor usage is not the driving force for the emergence of omicron variants. Instead, achieving the immune evasion abilities appears to be the selection pressure behind the emergence of omicron variants.^28–30^ Our binding affinity data clearly show that all the omicron variants attain class I and class II antibodies escape abilities by incorporating mutations at the hotspot residues for each specific class of antibody recognition. K417N and Y505H point mutation significantly reduce class I antibody binding affinities, while E484A appears critical for escaping class II antibody neutralization. E484A and E484A/Q493R double mutations cause a 33-38% reduction in binding affinity for the approved therapeutic monoclonal antibodies, Bamlanivimab (LY-CoV555). The Q498R RBD mutation observed across all the omicron variants can reduce ~12% binding affinity for REGN10987, a class III therapeutic antibody approved in combination with REGN10933 for emergency approval. The L452R/Q498R double mutation causes a ~6% decrease in binding affinities for another class III therapeutic antibody, LY-CoV1404 (Eli Lilly’s IgG1 MAbs). Our data delineate the effect of critical point mutations on the observed immune evasion abilities of currently circulating omicron variants.

## Data availability statement

The data that support the findings of this study are available from thecorresponding author upon resonable request.

## Acknowledgement

This work is supported by the COVID-19 HPC Consortium (Grant ID:CHE210070) and authors gratefully acknowledge Microsoft AI for Health for providing technology support to carry out the research.

